# MAPK11 (p38β) is a major determinant of cellular radiosensitivity by enhancing IR-associated senescence

**DOI:** 10.1101/2022.09.12.506954

**Authors:** DM Fernández-Aroca, N García-Flores, S Frost, J Jiménez-Suarez, A Rodríguez-González, P Fernández-Aroca, S Sabater, I Andrés, C Garnés-García, B Belandia, FJ Cimas, D Villar, MJ Ruiz-Hidalgo, R Sánchez-Prieto

## Abstract

**Background and purpose:** MAPKs are among the most relevant signalling pathways involved in coordinating cell responses to different stimuli. This group includes p38MAPKs, constituted by 4 different proteins with a high sequence homology: MAPK14 (p38α), MAPK11 (p38β), MAPK12 (p38γ) and MAPK13 (p38δ). Despite their high similarity, each member shows unique expression patterns and even exclusive functions. Thus, analysing protein-specific functions of MAPK members is necessary to unequivocally uncover the roles of this signalling pathway. Here, we investigate the possible role of MAPK11 in the cell response to ionizing radiation (IR).

**Materials and methods:** We developed MAPK11/14 knockdown through shRNA and CRISPR interference gene perturbation approaches, and analysed the downstream effects on cell responses to ionizing radiation in A549, HCT-116 and MCF-7 cancer cell lines. Specifically, we assessed IR toxicity by clonogenic assays; DNA damage response activity by immunocytochemistry; apoptosis and cell cycle by flow cytometry (Annexin V and propidium iodide, respectively); DNA repair by comet assay; and senescence induction by both X-Gal staining and gene expression of senescence-associated genes by RT-qPCR.

**Results:** Our findings demonstrate a critical role of MAPK11 in the cellular response to IR by controlling the associated senescent phenotype, and without observable effects on DDR, apoptosis, cell cycle or DNA damage repair.

**Conclusion:** Our results highlight MAPK11 as a novel mediator of the cellular response to ionising radiation through the control exerted onto IR-associated senescence.

**Highlights:**

- Genetic perturbation of MAPK11, but not MAPK14, promotes radiosensitivity in a panel of tumor cell lines.
- Abrogation of MAPK11 did not modify DNA damage response, proliferation, apoptosis or cell cycle in response to ionizing radiation
- MAPK11 controls ionizing radiation-induced senescence
- MAPK11 expression could be a novel target and biomarker for radiosensitivity

## Introduction

Radiotherapy, applied to around 50% of cancer patients, has become a cornerstone in cancer therapy [1], being especially relevant in some types of tumours such as breast, colon or lung [2]. It is therefore essential to uncover the molecular and biological processes triggered by ionising radiation (IR) which could improve the effectiveness of treatments. Consequently, the search of mechanisms responsible for sensitisation and resistance to radiotherapy, both *de novo* and acquired, has been a long-standing issue in radiobiology [3–6]. Several signalling pathways [7], biological processes [8,9], genetic alterations [10,11], and even epigenetic modifications [12] have been related to the cellular response to IR. However, we still do not have a complete picture of the molecular elements involved in this biological response that could contribute to improve and personalise radiotherapy. Within p38MAPK family, four proteins can be found: MAPK14, MAPK11, MAPK12 and MAPK13. With the term p38MAPK we will refer to all four proteins. Despite these members sharing high sequence homology [13], each of them shows not just different tissue expression patterns [14,15] but also specific functions in different biological processes [16,17] and different implications in cancer [18]. Furthermore, even opposite roles for each member have been described [19] (e.g. in activating AP-1-dependent transcription in breast cancer cell lines [20] or in pancreatic cancer [15,21]). Most of the current evidence linking p38MAPK and cancer focuses onto MAPK14, due to its ubiquitous and abundant expression [22]. However, a growing body of data support a key role in cancer for other members of the family, for instance MAPK12 and MAPK13 [23,24]. Indeed, recent evidence indicates important roles for MAPK11 in cancer and its therapy (for a review see [25]). Nonetheless, p38MAPK signalling pathway has been linked with the response to DNA damage and, specifically, to IR. TAO kinases are able to activate p38MAPK through ATM/ATR pathways in response to IR [26,27], emerging as regulators of p38-mediated response to DNA damage [26]. It is also remarkable that p38MAPK has been found to be critical in biological/biochemical processes triggered by IR, such as IR-induced apoptosis [28] or AKT activation [29]. Among the p38MAPK members which have been specifically proposed as mediators of cell response to IR, MAPK14 has been implicated in a plethora of effects ranging from autophagy [30] up to cell cycle control [31], whereas the rest of members (MAPK11, 12 13) has been barely studied in response to IR [32–34]. In addition, it is noteworthy that the vast majority of publications assessing p38MAPK roles are based on pharmacological approaches including those related to IR (e.g. [35–42]), which at best allow to distinguish MAPK11/14 from MAPK12/13 not addressing protein-specific functions for each p38MAPK family member.

Against this background, we aimed to clarify the specific role of MAPK11 in the cellular response to radiotherapy in different experimental models, including colon, lung, and breast cancer cell lines by using genetic approaches to fully exploit the potential role of this particular MAPK in radiotherapy.

## Materials and Methods

### Cell lines and plasmids

A549 (Lung cancer), HCT-116 (Colon cancer) MCF-7 (Breast cancer) and HEK293T have been cultured as previously described [43]. Cells were maintained in 5% CO_2_ and 37°C; and grown in Dulbecco’s modified Eagle’s medium supplemented with 10% fetal bovine serum, 1% glutamine and 1% Penicillin/Streptomycin. All cell culture reagents were provided by Lonza.

Plasmids used for shRNA interference (Sigma-Aldrich) were as follows: Human pLKO.1-puro-shRNAMAPK14 (Sigma SHCLNG-NM_001315; TRCN0000000511), Human pLKO.1-puro-shRNAMAPK11 (Sigma SHCLNG-NM_002751; (TRCN0000199694), and pLKO.1-puro empty vector (Sigma SHC001). For CRISPR interference (CRISPRi) were as follows: TRE-dCas9-KRAB-IRES-GFP (Addgene #85556) [44], pU6-sgRNA-puro-BFP (Addgene #60955) [45].

### Transfections and infections

Lentiviral production and cell infection were performed as previously described [46,47].

### Inducible CRISPR interference (CRISPRi)

A549 cells were infected with lentiviruses containing TRE-dCas9-KRAB-IRES-GFP [48], treated for 4 days with 1 μg/ml doxycycline and then GFP-positive cells were sorted by flow cytometry. Next, dCas9-expressing cells were infected with pU6 plasmids harbouring non-target control (NTC) or sgMAPK11 gRNAs and selected with 1μg/ml puromycin for 3 days. To achieve full dCas9 expression, cells were treated with 1 μg/ml doxycycline 5 days prior to cell seeding and maintained over the course of the experiments.

gRNAs (Supplementary Table 1) were designed with CHOPCHOP [49], purchased from IDT, and cloned into pU6-sgRNA-puro-BFP as previously described [50].

### Western Blotting

Protein quantification and western blotting was performed as previously described [33]. Antibodies used are summarized in Supplementary Table 2. Images show a representative experiment out of three with similar results.

### Immunocytochemistry

Cells were grown onto SPL cell culture slides (Labclinic) 24 h prior to irradiation. After treatment cells were fixed, permeabilized and incubated with the indicated antibodies (Supplementary Table 2) as previously described [51]. Positive immuno-fluorescence was detected using a Zeiss Apotome fluorescence microscope and processed using Zen 2009 Light Edition program (Zeiss). Foci quantification was performed with CellProfiler (Broad) [52]. Images show a representative cell from a minimum of 100 quantified (5 fields per sample captured). Data shown are the average of, at least, three independent experiments.

### RNA isolation, reverse transcription and Real-time Quantitative PCR

Total RNA was obtained as previously described [53]. cDNA synthesis was performed with RevertAid First Strand cDNA synthesis Kit (Thermo Scientific) following manufacturer’s protocol in an iCycler thermal cycler (Biorad). Real time PCR was performed with Fast SYBR Green Master kit (Thermo Scientific) in a 7500 Fast Real-Time PCR instrument (Applied Byosystems). PCR conditions were as previously described [53]. Primers for all target sequences were designed by using NCBI BLAST software and purchased from Merck as DNA oligos. Primer sequences can be found in Supplementary Table 1. Data shown are the average of, at least, three independent experiments performed in triplicate.

### Irradiation and clonogenic assays

Cells were irradiated by the technical staff of Radiotherapy Unit at University General Hospital of Albacete, in a Clinac Low Energy 600C linear electron accelerator from Varian (Palo Alto, California, USA) at a dose rate of 600 cGy/min in a radiation field of 40×40 cm. Clonogenic assays were performed and valuated as previously described [53,54]. Plates were photographed and colonies were counted with the ImageJ plugin “Cell counter”. Colonies with less than 5 mm diameter were discarded. Values were referred to unirradiated controls, set at 1. SF2Gy was calculated by applying a linear-quadratic model [55]. Data shown are the average of, at least, three independent experiments performed in triplicated cultures.

### β-galactosidase activity

Six days after irradiation, cells were washed in PBS, fixed for 5 min (room temperature) in 12% formalin, washed twice for 5 minutes, and incubated for 16 h at 37°C (no CO_2_) with fresh SA-β-Gal staining [56]. Images were acquired at 10x using Zeiss Apotome. Images show a representative field out of 5 acquired per sample (minimum of 100 cells quantified per condition). Data shown are the average of three independent experiments.

### Flow cytometry

For cell cycle analysis, 10^5^ cells were seeded in 6 cm plates 24 hours prior to irradiation and cell cycle was analysed as previously described at indicated times [43]. For apoptosis detection, 10^5^ cells were seeded in 6 cm plates, 24 h later cells were treated with IR and after 48 h apoptosis was detected with Annexin V-FITC (Immunostep) following manufacturer’s instructions.

Samples were processed in a MACSQuant Analyzer 10 (Miltenyi Biotec). Data were analysed by using FlowingSoftware (University of Turku). Data shown are the average of, at least, three independent experiments performed.

### Comet Assay

DNA fragmentation and repair was measured with the alkaline comet assay [57]. Images were acquired at 10x magnification using Zeiss Apotome fluorescence microscope and analysed with the plugin OpenComet [58] (ImageJ) to measure tail moment (DNA% in tail * tail length). Data shown are the average of three independent experiments.

### Cell proliferation measurements

For cell proliferation measurements, 10^4^ cells/well were seeded in 24-well plates and proliferation was analysed at 1, 2 and 3 days later by an MTT-based assay as previously described [33]. Data shown are the average of three independent experiments performed in triplicated cultures.

### Statistical analysis

Data are presented as mean ± standard deviation (S.D). Statistical significance was evaluated by Student’s t test or ANOVA using GraphPad Prism v9.0 software. The statistical significance of differences is indicated in figures by asterisks as follows: *p < 0.05, **p < 0.01 and ***p<0.001.

For Kaplan-Meyer curves, analysis was performed in cBioportal [59] by using curated TCGA Breast Cancer dataset. The differences between survival curves were examined using the log-rank test. Patients were segregated depending on MAPK11 mRNA levels.

## Results

Given the evidence of p38MAPK implication in the cellular response to IR, we first aimed to interrogate the specific role of MAPK11 and MAPK14, the two highly and ubiquitously expressed members of p38MAPK across A549 (lung), HCT-116 (colon) and MCF-7 (breast) cancer cells lines. After achieving an effective MAPK14 or MAPK11 knockdown by shRNA (Fig. 1A, D, G and Sup. Fig. 1), we analysed viability after IR exposure by 15-days clonogenic assays. We did not observe a significant effect upon MAPK14 abrogation in any of the cell lines tested, however, knockdown of MAPK11 led to a significant reduction in cell survival after IR exposure (Fig. 1B, E, H)) in all three cell lines, but did not affect proliferation in the absence of radiation (Sup. Fig. 2). Indeed, SF2Gy showed a marked and specific decrease upon MAPK11 ablation (Fig. 1C, F, I), suggesting a potential role for this MAPK in radiobiology.

**Fig. 1.**
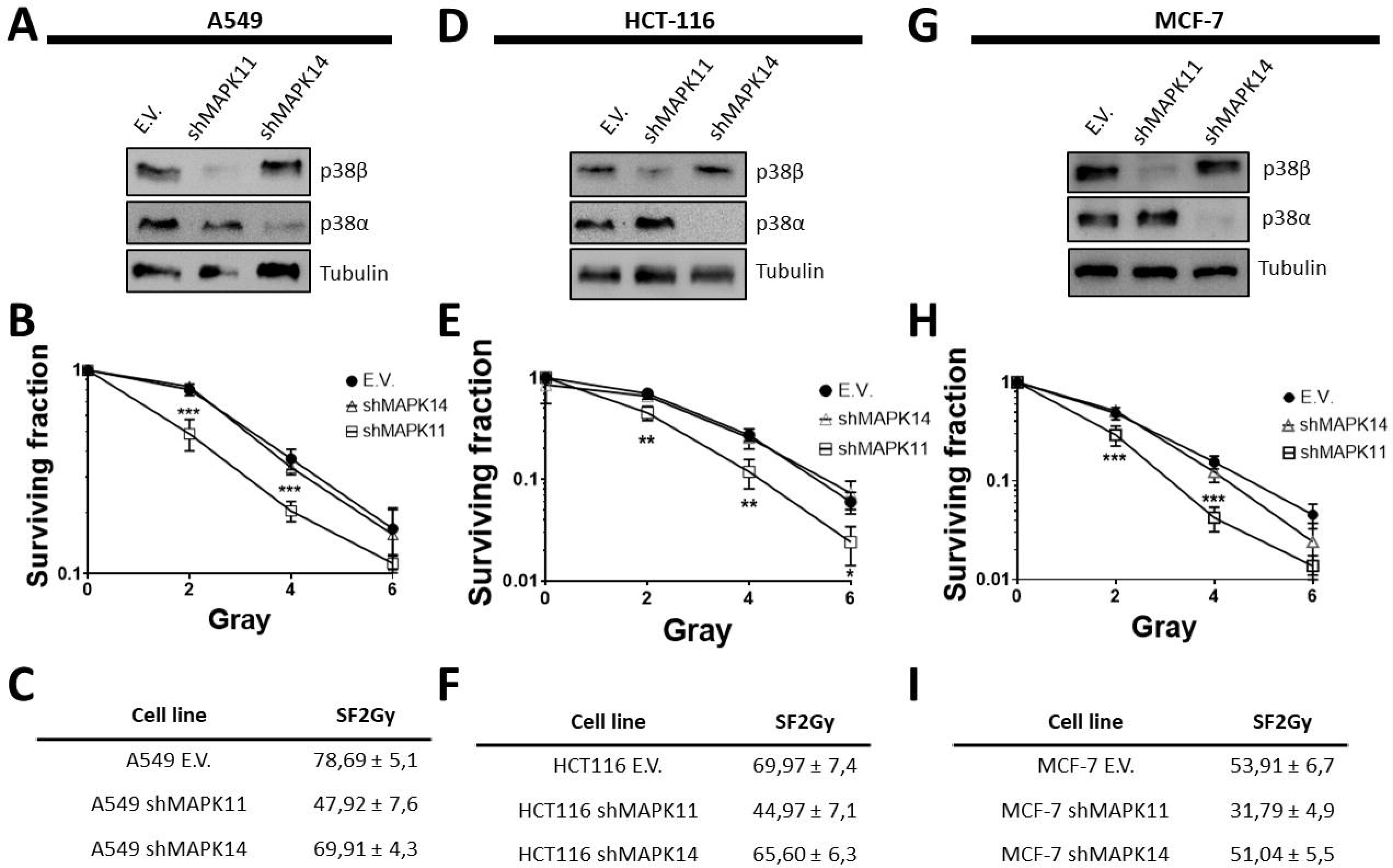
Genetic abrogation of MAPK11 promotes radiosensitivity in A549, HCT-116 and MCF-7 cell lines. A) A549 cells were infected with lentiviruses carrying empty vector (E.V), shRNA for MAPK11 (shMAPK11) or MAPK14 (shMAPK14). Genetic interference was evaluated by western blot using tubulin as a loading control. Bars denote standard deviation (S.D). B) Clonogenic assays for A549 cells infected with E.V, shMAPK11 or shMAPK14 and exposed to the indicated doses of X rays. Surviving fraction was normalized to respective unirradiated controls. Bars mean standard deviation (S.D). C) A549 E.V., shMAPK11 and shMAPK14 surviving fraction at 2 Gy (SF2Gy) ± S.D. calculated by lineal-quadratic model. D) Same as in A) for HCT-116 cells. E) Same as in B) for HCT-116 cells. F) Same as in C) for HCT-116 cells. G) Same as in A) for MCF-7 cells. H) Same as in B) for MCF-7 cells. I) Same as in C) for MCF-7 cells.

Since the activity of DNA Damage Response (DDR) is one of the most important cell pathways activated in response to IR, we next investigated whether MAPK11 plays any role in this signalling pathway by choosing as experimental model the A549 cell line. To this end, we analysed activation of key molecules in the cellular response to DNA damage, such as ATM and homologous-recombination repair mediator BRCA1. These experiments showed no differences in the number of pATM/pBRCA1 foci per nuclei in cells with reduced MAPK11 expression compared to control cells (E.V.) (Fig. 2A). This result was also confirmed by comet assay, in which no differences in DNA repair capacity were found (Fig 2. B). Furthermore, we confirmed these results in MCF-7 and HCT-116 cell lines (Sup. Fig. 3), thus concluding MAPK11 does not play a direct role in mediating DDR activity.

**Fig. 2.**
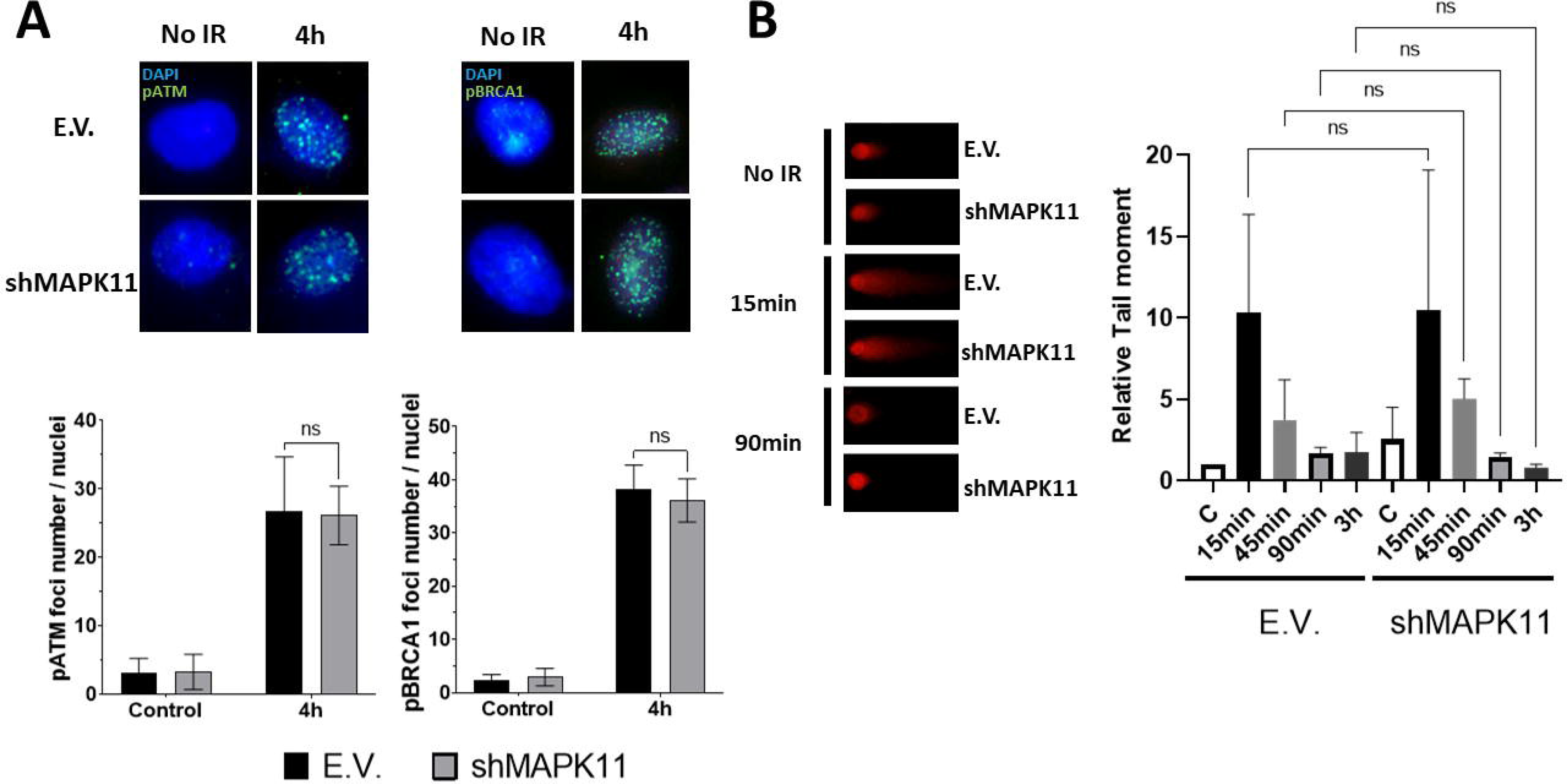
MAPK11 does not modulate DDR activity and DNA repair in response to ionising radiation. A) *Upper panel*: A549 cells harbouring empty vector (E.V.) or shMAPK11 were plated onto cell culture slides 24 h prior to irradiation (10 Gy) and 4 h later cells were fixed and processed for immunocytochemistry against phospho-ATM (Ser1981) or phospho-BRCA1 (Ser1524). Images show a representative cell out of a minimum of 100 analysed. *Lower panel*: Quantification of phospho-ATM or phospho-BRCA1 foci number per nuclei in three independent experiments. Bars mean standard deviation (S.D). B) *Left panel*: Comet assay analysis of A549 E.V. and shMAPK11 cells. After irradiation (10Gy) DNA damage was evaluated at the indicated times. Images show a representative cell out of a minimum of 100 analysed. *Right panel*: Histogram showing comet tail moment normalized to E.V. unirradiated controls. Bars mean standard deviation (S.D).

Next, we studied MAPK11 effects in terms of IR-associated cell cycle blockage and apoptosis induction, as these are other two relevant, early cellular responses to DNA damage. We did not observe MAPK11 to have a determinant role onto G2/M cell cycle phase accumulation 24 h after treatment with radiotherapy in A549 cells (Fig. 3A). Interestingly, we observed a slight but non-significant premature release from G2/M blockage in shMAPK11 cells 48 h after IR. Regarding IR-induced apoptosis, we did not find significant differences in induction of apoptosis 48h after IR exposure in A549 (Fig 3B), HCT-116 or MCF-7 cell lines (Sup. Fig. 3). Therefore, the collective evidence from these experiments discards a direct role of MAPK11 in the early cellular response to IR, at least within the first 48 h after induction of DNA damage.

**Fig. 3.**
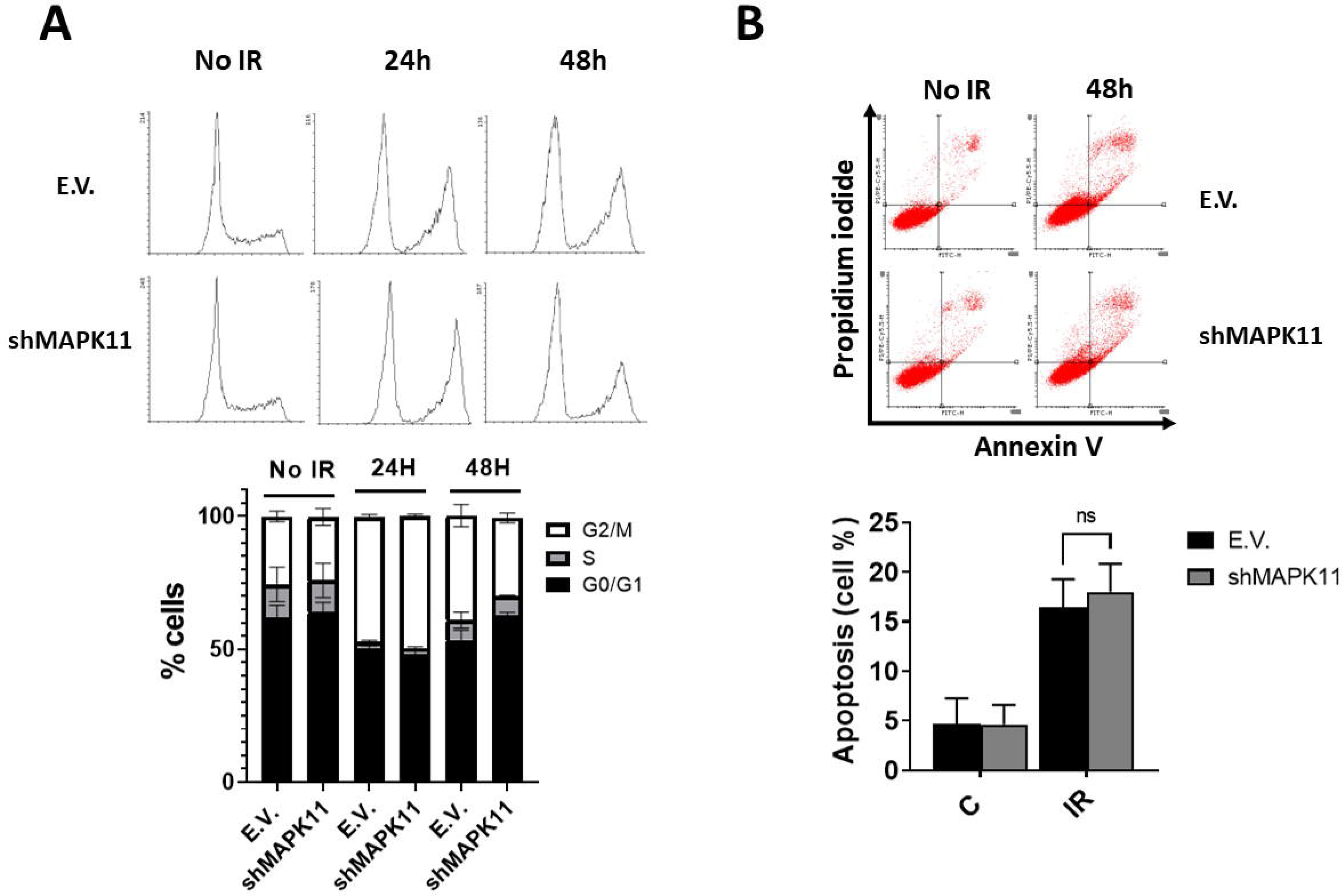
MAPK11 does not deregulate cell cycle and apoptosis after irradiation in A549 cell line. A) *Upper panel*: Image of a representative cell cycle profile in A549 E.V. and shMAPK11 cells irradiated at 10 Gy. Cell cycle was evaluated by flow cytometry at indicated times after IR. *Lower panel*: Histogram showing the average of three independent experiments representing the percentage of population in the different phases of the cell cycle. Bars mean S.D. B) *Upper panel*: Graphical representation of apoptosis induction in A549 E.V. and shMAPK11 cells 48 h after irradiation (10 Gy) by staining with Annexin V-FITC/Propidium Iodide for assay by flow cytometry. *Lower panel*: Histogram showing the average of three independent experiments to evaluate the percentage of apoptotic A549 E.V. or shMAPK11 cells 48 h after IR (10 Gy). Bars mean S.D.

In light of the lack of effect in early responses, we reasoned out the radiosensitivity we observed in 15-days clonogenic assays (Fig 1) could be triggered by a later cellular response. In order to study other biological consequences of IR, we investigated induction of cellular senescence, which is known to onset several days after irradiation [60]. To this end, we assessed IR-induced β-Gal activity 6 days after IR in cells infected with E.V. or shMAPK11, as a well-stablished marker of senescence. As shown in Fig. 4A, A549 cells harbouring shRNA targeting MAPK11 undergo enhanced induction of senescence-like phenotypes compared to E.V. cells. In addition, we confirmed these results by analysing gene expression of well-stablished senescence markers: IL-1β, p21, IL-6 and IL-8 [61], observing a significant induction for all of them after MAPK11 knockdown compared to E.V. cells (Fig. 4B). In sum, our results indicate that the lack of MAPK11 could promote a marked increase in cellular senescence secondary to irradiation.

**Fig. 4.**
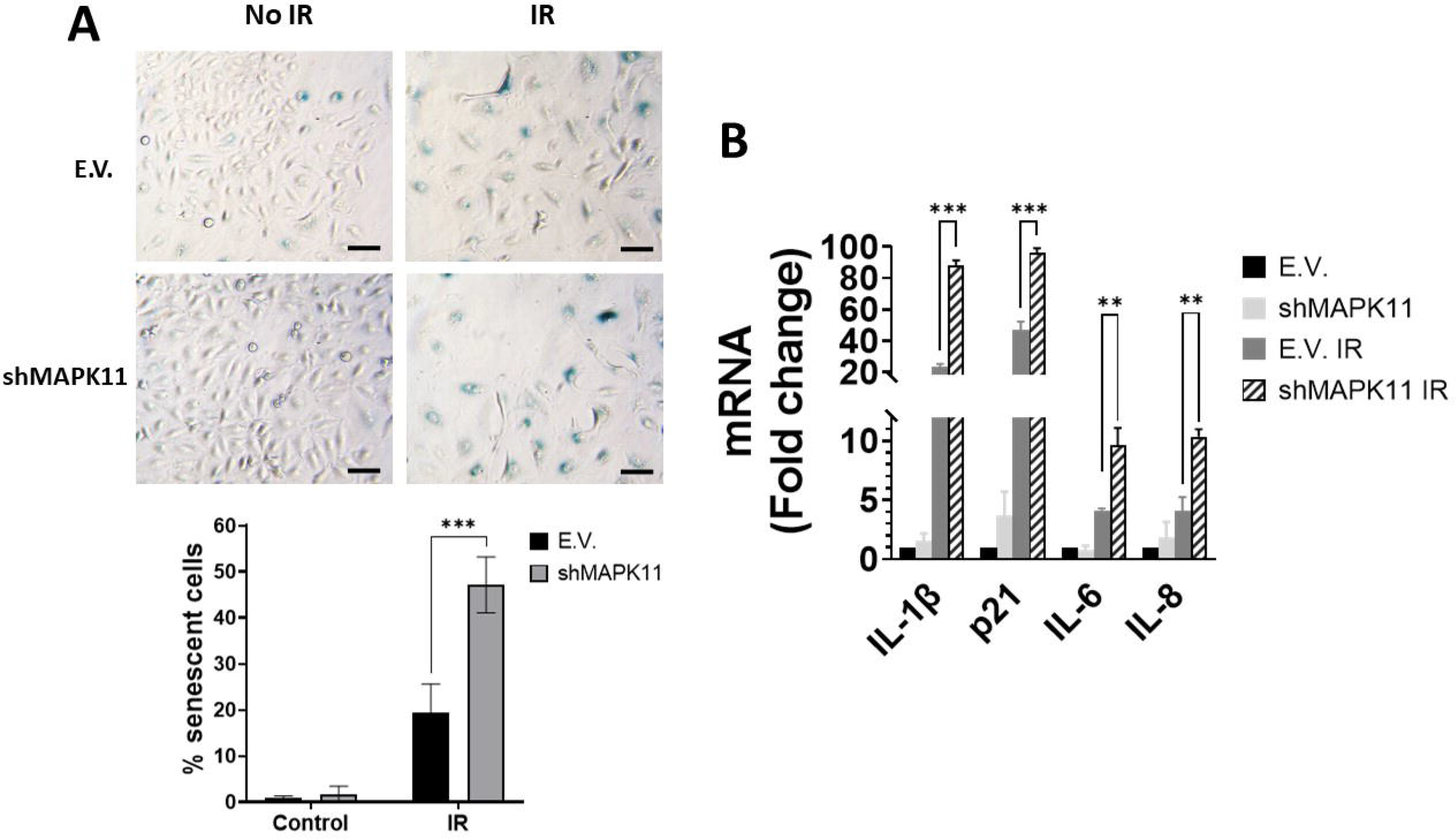
MAPK11 genetic interference enhances cell senescence in response to ionizing radiation. A) Upper panel: A549 E.V. or shMAPK11 cells were irradiated (10 Gy) and 5 days later β-Gal activity was detected by X-gal staining. A representative image is shown. Scale bars represent 10 μm. Lower panel: Histogram showing the average of, at least, three independent experiments representing the percentage of positive senescent cells. Bars mean S.D. B) Gene expression of indicated senescence-associated genes was evaluated 5 days after IR (10 Gy) in A549 E.V. or shMAPK11 cells by RT-qPCR using GAPDH as endogenous control. Data were referred to unirradiated E.V. cells. Bars mean S.D.

To verify our observations based on shRNA, we developed a complementary epigenetic perturbation approach based on doxycycline(dox)-inducible CRISPR interference. After achieving an effective knockdown (Fig 5A, Sup 4A), we performed clonogenic assays with A549 cells harbouring a dox-inducible dCas9-KRAB and a non-target control gRNA (NTC) or a gRNA targeting MAPK11 promoter (sgMAPK11). In line with shRNA data, CRISPRi knockdown of MAPK11 was able to sensitize A549 cell line to IR (Fig 5B) showing a lower SF2Gy (NTC=76.05±8.6; sgMAPK11=43.65±3.5). Moreover, both biochemical and biological effects of CRISPRi knockdown were similar to those obtained with shRNA: no effect was observed on DNA damage repair, p-ATM foci formation and apoptosis induction (Sup. Fig. 4); while, as expected, senescence induction was increased in cells upon MAPK11 CRISPRi knockdown, in terms of both β-Gal staining and expression of senescence-related genes (Fig 5C, D). Collectively, these results confirm those obtained with shRNA interference, and strongly suggests an important role for MAPK11 in mediating the cellular response to IR.

**Fig. 5.**
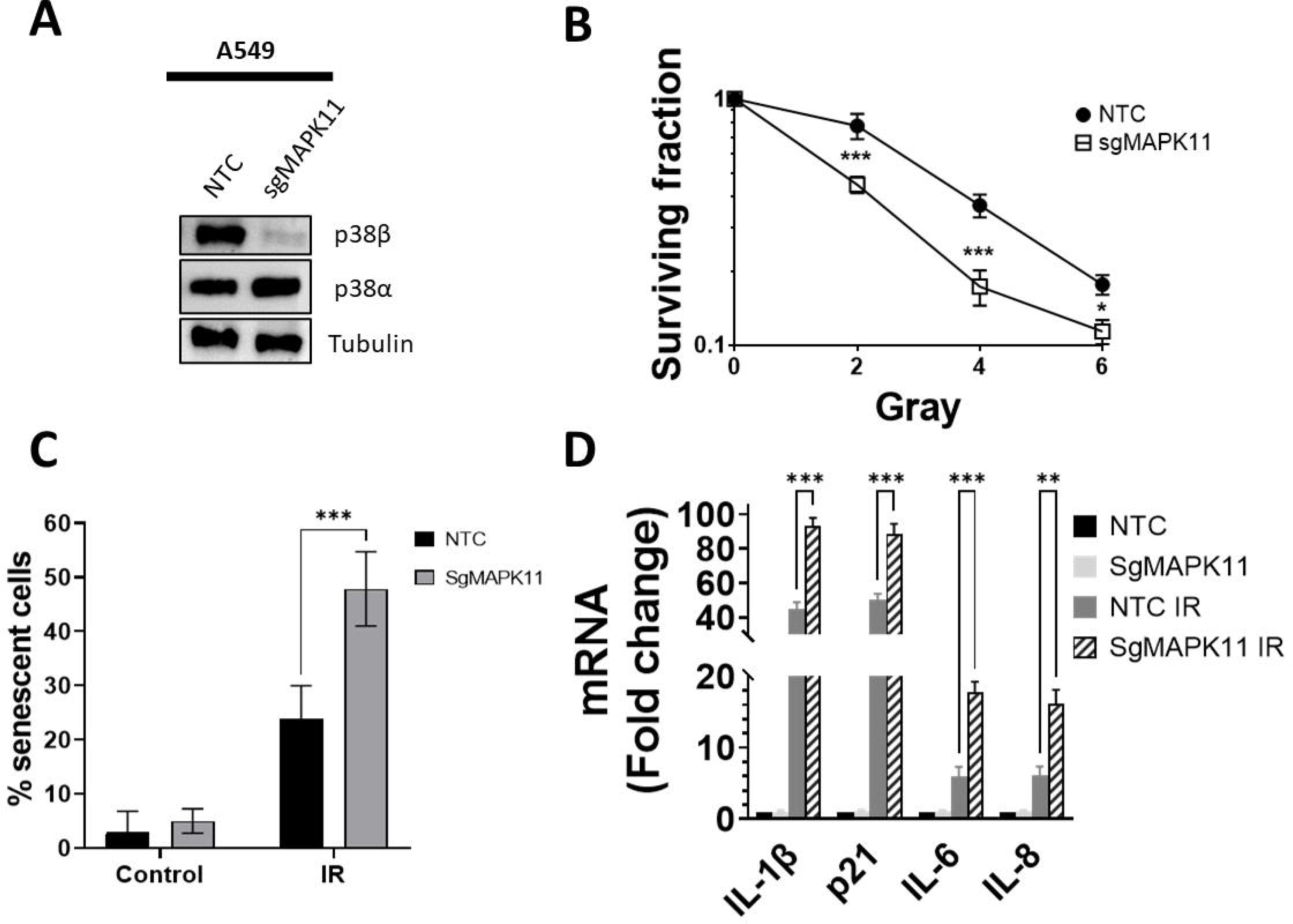
Epigenetic perturbation of MAPK11 by CRISPRi confirms its role in cell response to ionizing radiation. A) A549 cells expressing dCas9 were infected with lentiviruses carrying non-target-control (NTC) or MAPK11-targeted sgRNAs (sgMAPK11). Interference was evaluated by RT-qPCR using GAPDH as an endogenous control (right) and by western blot using tubulin as a loading control (left). Bars mean standard deviation (S.D). B) Clonogenic assays for A549 NTC and sgMAPK11 cell exposed to the indicated doses of X rays. Bars mean standard deviation (S.D). C) Histograms show the average of three independent experiments representing the percentage of positive senescent cells evaluated by X-Gal staining 5 days after IR (10 Gy). Bars indicate S.D. D) Gene expression of indicated senescence-associated genes was evaluated 5 days after IR (10 Gy) in A549 NTC or sgMAPK11 cells by RT-qPCR using GAPDH as endogenous control. Data were referred to unirradiated NTC cells. Bars mean S.D.

To evaluate the generality of our observations, we assessed β-Gal activity and induction of IL-1β, IL-6, IL-8 and p21 in response to IR in HCT-116 and MCF-7 cell lines with and without MAPK11 expression. In both experimental models, we observed an enhancement of β-Gal activity and gene expression profiler associated to IR dependent senescence (Fig 6A; B, C and D); thus confirming MAPK11 could be involved in the induction of senescence in response to IR. Finally, to evaluate the clinical implications of our findings, we performed an *in silico* analysis by using cBioportal platform, which stores information from the TCGA database including patient data upon radiotherapy treatment for almost all tumour types. The limited number of patients treated with radiotherapy for most tumour types restricted our analyses to breast cancer (643 patients, Fig 6E)). Nevertheless, this *in silico* analysis points to a clear relationship between MAPK11 expression levels and clinical outcome, in terms of overall survival, of patients undergoing radiotherapy treatment, thus supporting a key role for this MAPK in radiobiology and radiotherapy.

**Figure 6.**
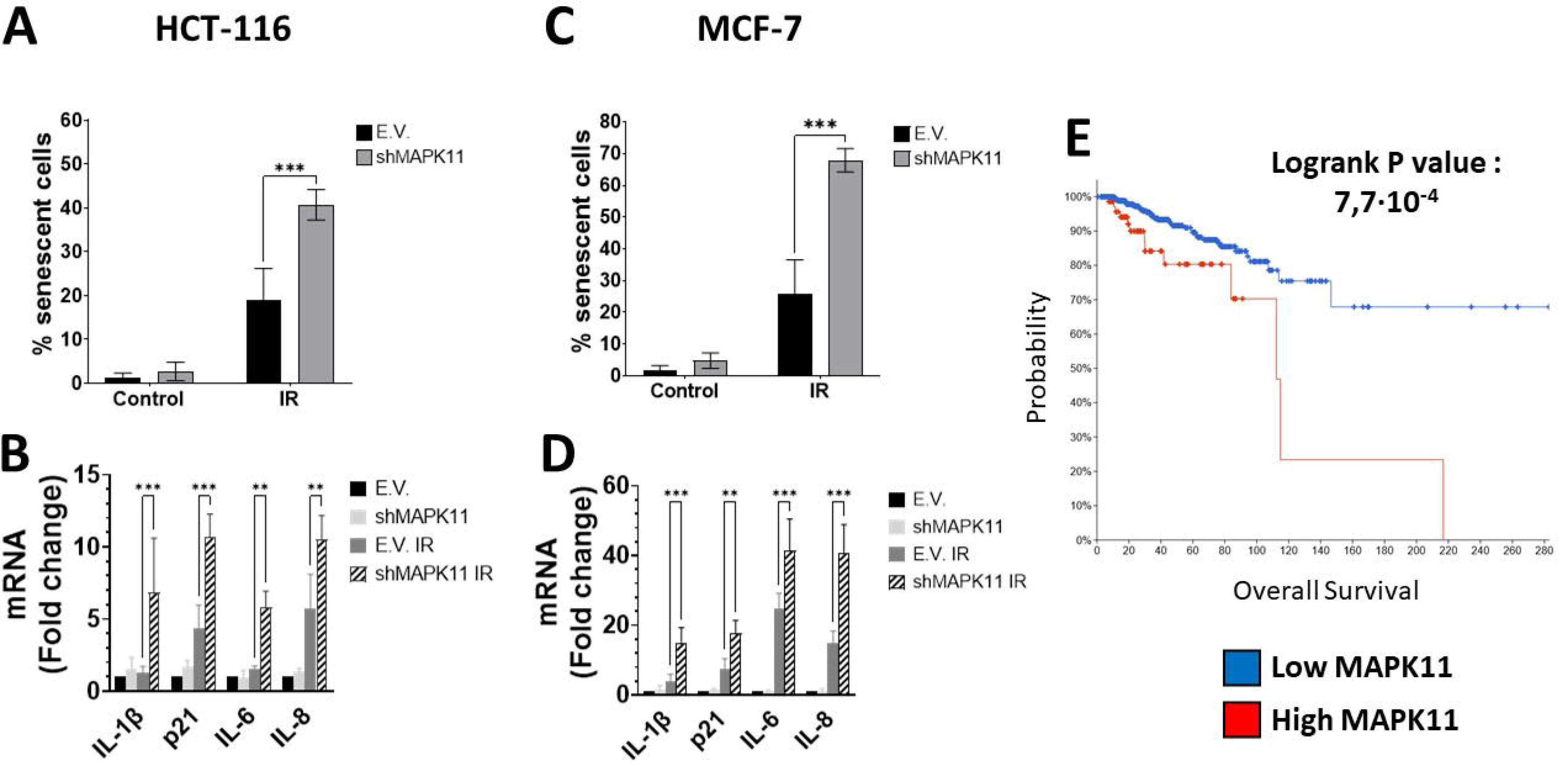
Genetic interference of MAPK11 increases induction of senescence in response to ionizing radiation in HCT-116 and MCF-7 cell lines. A) HCT-116 E.V. and shMAPK11 cells were irradiated (10 Gy) and 5 days later senescence was evaluated by X-Gal staining. Histogram shows the average of three independent experiments representing the percentage of positive senescent cells. Bars mean S.D. B) Gene expression of indicated senescence-associated genes was evaluated 5 days after IR (10 Gy) in HCT-116 E.V. and shMAPK11 cells by RT-qPCR using GAPDH as endogenous control. Data were referred to unirradiated E.V. cells. Bars mean S.D. C) Same as in A) for MCF-7 cells. D) Same as in B) for MCF-7 cells. E) Kaplan–Meyer comparing prognosis in terms of Overall Survival (OS) for two groups of patients, those with high (mRNA expression > 1.2 S.D., n=57) and low (mRNA expression < 1.2 S.D., n=491) expression levels of MAPK11 for IR-treated breast patients from TCGA dataset.

## Discussion

IR triggers a broad, complex, and highly regulated cellular response involving a wide variety of signalling pathways including the superfamily of MAPKs. Despite p38MAPK, mainly MAPK14, has been deeply studied in relation to IR, no connection has been established, to the best of our knowledge, between MAPK11 and the cellular response to IR. In the present work we have observed how MAPK11 abrogation drives a marked increase in cellular sensitivity to IR, regardless of the genetic perturbation approach applied and the cancer cell line studied. Interestingly no differences in DNA damage response, cell cycle arrest and apoptosis, all, of then early responses to IR [62], were detected in association with those observed in clonogenic assays. Conversely, radiosensitisation associated with MAPK11 abrogation could be explained by a promotion of IR-induced senescence, a critical late-onset cellular effect associated to IR [60], establishing MAPK11 as a new key player in the cellular response to IR.

The key finding of our work is the observation of a clear enhancement of IR-associated senescence in the absence of MAPK11, with no apparent role for MAPK14. Although it has been reported that activation of p38MAPK, mainly MAPK14, can promote senescence associated to IR [63,64], this effect seems to be not applicable to several experimental systems in which senescence is promoted after inhibition of p38MAPK [65,66]. Moreover, all these previous works are based on the use of SB203580 inhibitor [67], which did not allow to distinguish between MAPK11 and 14; and in most, if not all the cases, MAPK14 was assumed to be the key mediator. Albeit no previous study has focused on MAPK11 and radiotherapy, several works have shown a role for MAPK11 in the response to oxidative stress, controlling the cell fate by blocking processes like apoptosis, senescence or autophagy in different experimental models such as brain [68], muscle [69] and cardiomyocytes [70]. Furthermore, it has been reported how abrogation of MAPK11 is required for the activity of tumor suppressor genes and some miRNAs, supporting an oncogenic and pro-survival role for this MAPK. [71,72]. In addition, other possibilities should be considered. For example, the histone deacetylase HDAC3, which has been recently proposed as a key regulator of Senescence Associated Secretory Phenotype [73], is known to interact with MAPK11 [74], supressing the transcriptional activity of ATF-2. Interestingly, ATF-2 is known to promote survival and efficient DNA repair after IR exposure [75] that could render radioresistance [76,77]. Furthermore, p38MAPK has been proposed to block senescence in response to IR by the control exerted onto miR-155 [78]. In sum, all this evidence supports that abrogation of MAPK11 promotes radiosensitivity. However, the molecular mechanisms by which MAPK11 induces senescence in response to IR and radiosensitivity needs to be further studied.

Finally, our observations could be relevant for the design of clinical trials in which p38MAPK inhibitors are combined with radiotherapy as in the case of glioblastoma [79]. The use of genetic approaches is an excellent proof of concept, yet their clinical implementation presents obvious difficulties relative to the use of pharmacological inhibitors., Hence, the search for specific molecules able to modulate MAPK11 (e.g. PROTAC technology [80])) is an interesting possibility that should be considered as a future research avenue. In addition, MAPK11 potential as a biomarker in response to IR would be worth exploring, especially in those tumors in which MAPK11 has been implicated and radiotherapy is a cornerstone of the treatment (e.g. female cancers or lung cancer [81,82])

In sum, our proof-of-concept results indicate that MAPK11 is a key player in the cellular response to IR trough the control of senescence. However, further research is necessary to fully exploit the potential of this MAPK as a target for radiosensitisation and/or as a predictive marker for the efficacy of radiotherapy.

## List of abbreviations

DDR: DNA damage response
DSB: Double strand break
gRNA: Guide RNA
IR: Ionizing radiation
MAPK: Mitogen Activated Protein Kinase
RT-qPCR: Quantitative real-time PCR
S.D.: Standard deviation
SF: Surviving fraction
shRNA: short hairpin RNA

## Acknowledgments

We would like to thank the staff from the Radiotherapy Unit at the Hospital General Universitario de Albacete for technical support. We also appreciate the technical assistance of SIB at CRIB and core facilities of IIB: We also appreciate the support of TALLER SOLIDARIO ARBOL DE LA VIDA (PERDOÑERAS), ASOCIACION COMARCAL CONTRA EL CANCER DE MOTILLA DEL PALANCAR and ACEPAIN in our research.

## Funding

This work has been supported with Grant PID2021-122222OB-I00 funded by MCIN/AEI/10.13039/501100011033, “ERDF A way of making Europe” to RSP. Also supported with funds from Fundación Leticia Castillejo Castillo (2021-AYUDA-32401) to RSP and MJRH. RSP and MJRH’s Research Institute and the work carried out in their laboratory, received partial support from the European Community through the FEDER. This work was also supported by a EMBO Short-Term Fellowship to DMFA. JJ holds a predoctoral research contract co-funded by the European Social Fund and UCLM. DV holds a personal Fellowship from the BHF (FS/18/39/33684). ARG and SF are supported by grants from Barts Charity and the BHF (MGU0501 and FS/18/39/33684 to DV). NGF has been supported by the Investigo Programme, within the framework of the Recovery, Transformation and Resilience Plan - financed by the European Union - Next Generation EU -, called by Order 190/2021, of 22 December, of the Regional Ministry of Economy, Business and Employment of the Regional Government of Castilla-La Mancha. FJC is funded by contracts for post-doctoral researchers for scientific excellence in the development of the Plan Propio I+D+i, co-financed by the European Social Fund Plus (ESF+).

## Conflicts of Interest

The authors declare that they have no known competing financial interests or personal relationships that could have appeared to influence the work reported in this paper.

## Notes

### Competing Interest Statement

The authors have declared no competing interest.

### Summary of Updates

We have slightly modified the manuscript structure and lenght to fit the journal's requirements, and reviewed spelling.

