## Supplementary figures and images for "MAPK11 (p38β) is a major determinant of cellular radiosensitivity by enhancing IR-associated senescence"

### Supplementary Figure 1

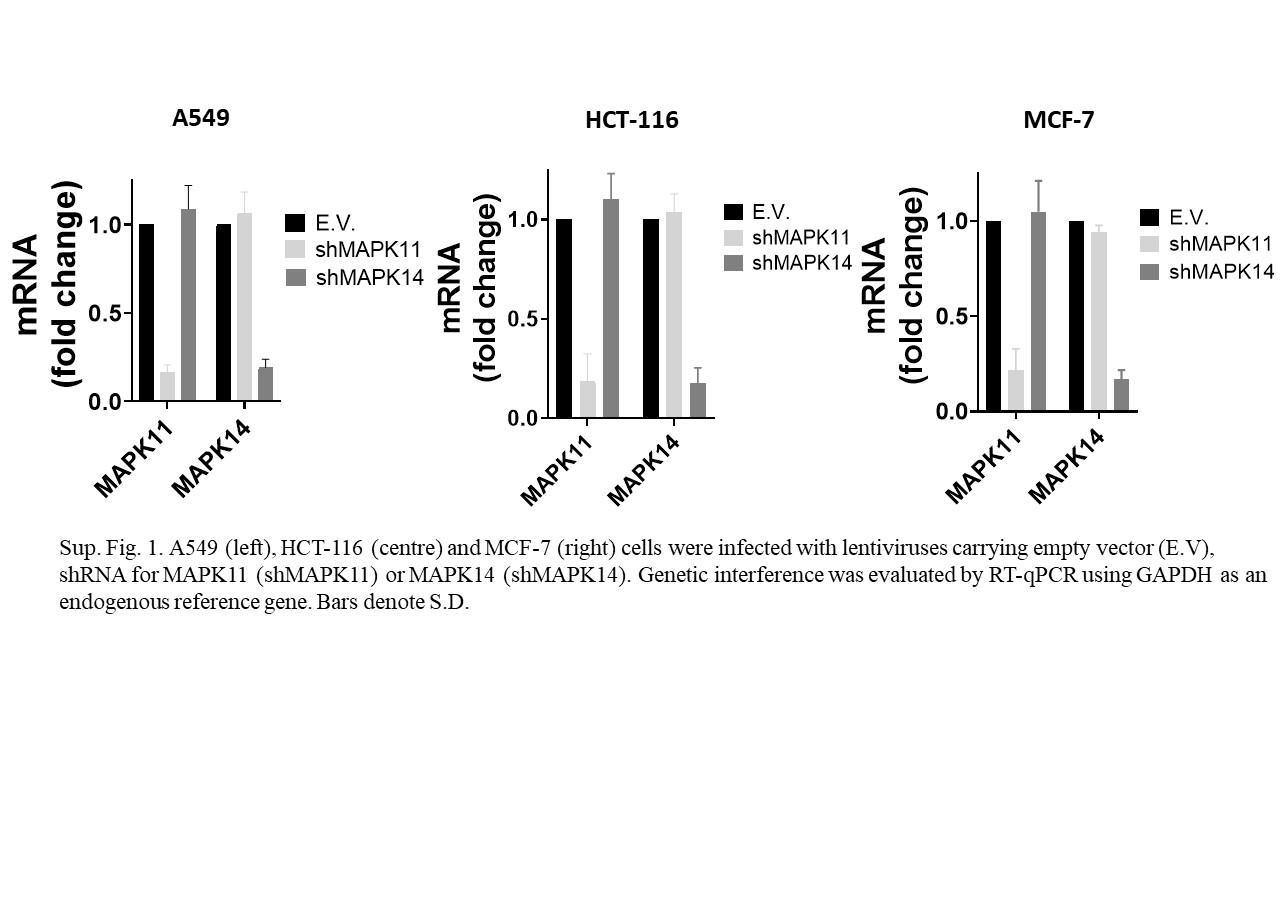

### Supplementary Figure 2

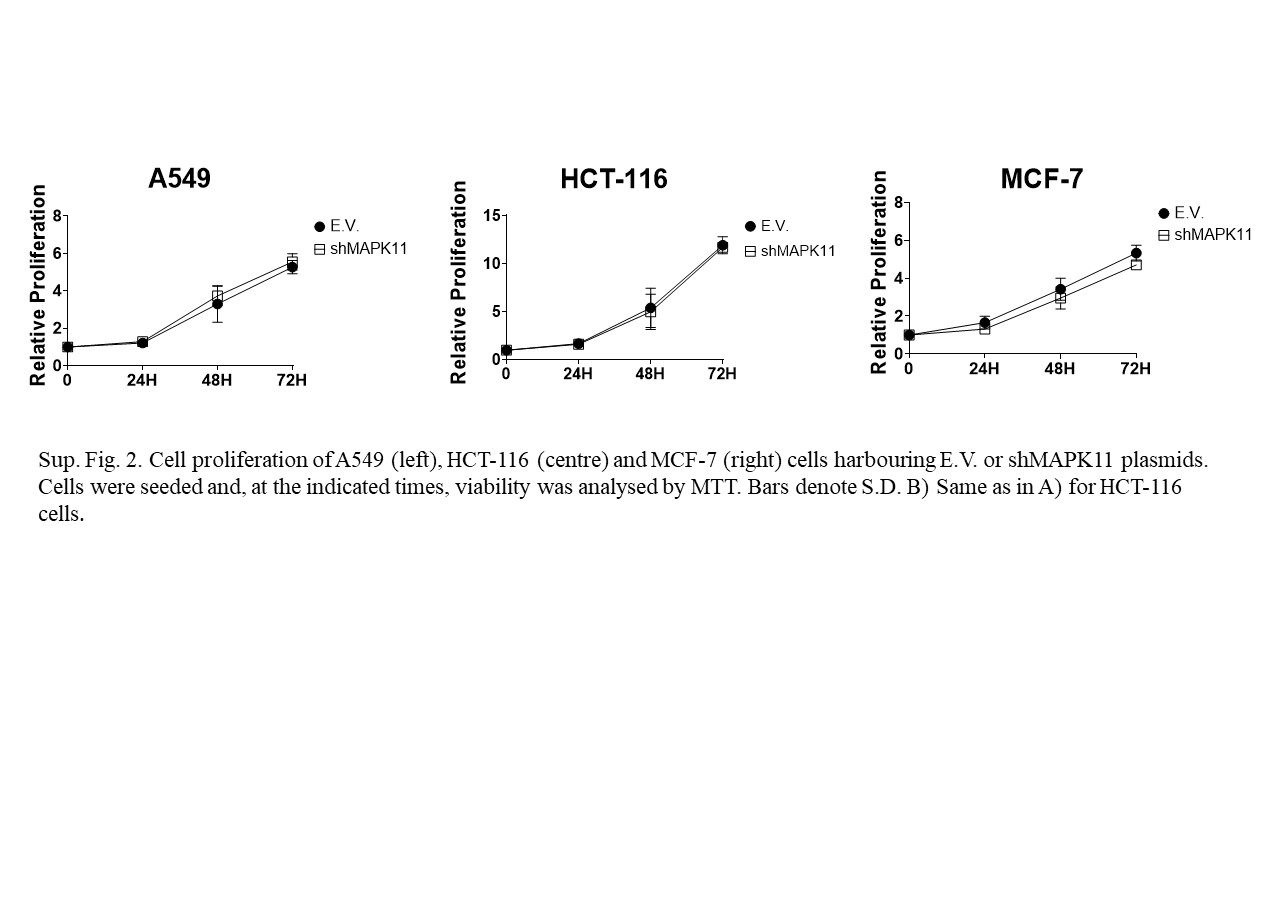

### Supplementary Figure 3

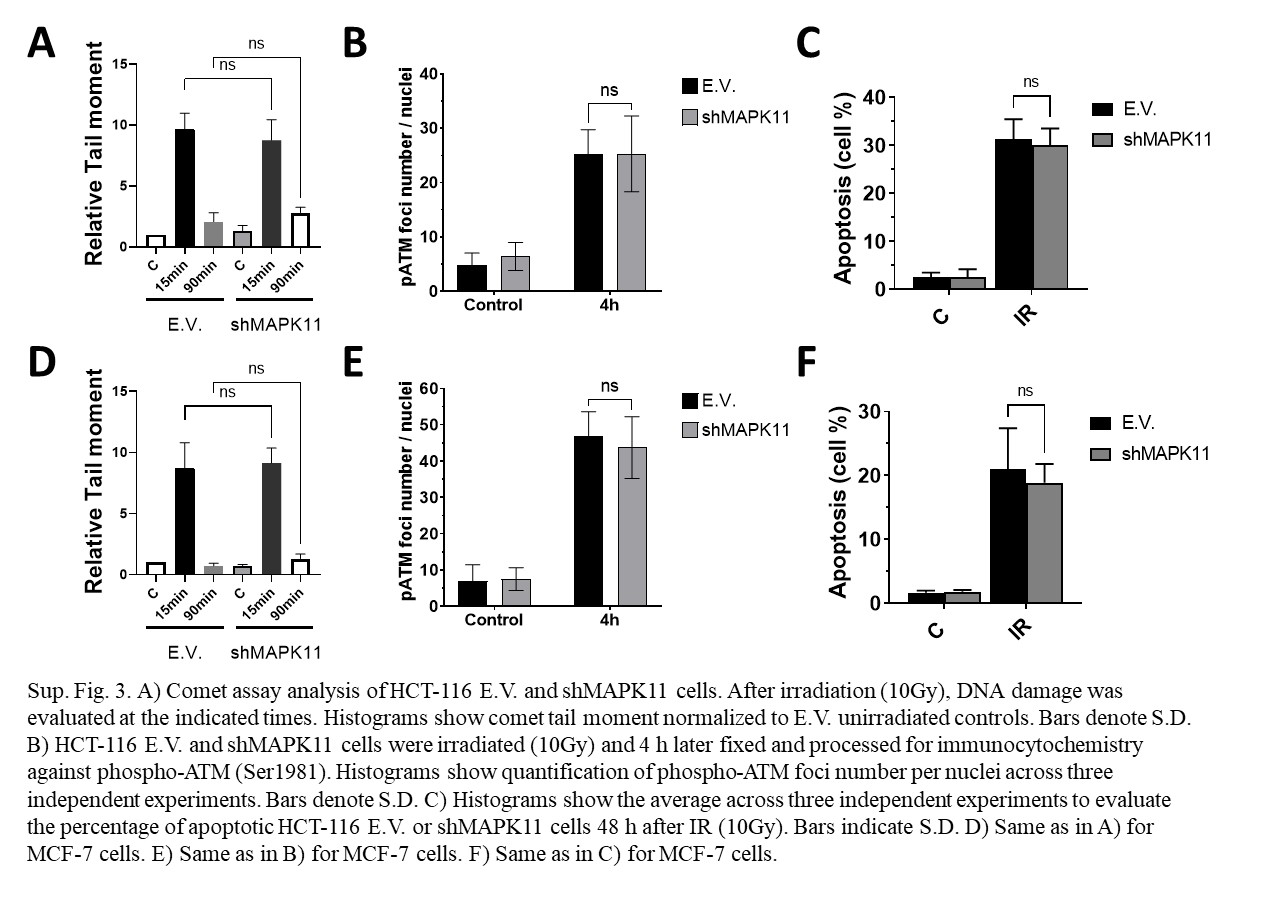

### Supplementary Figure 4

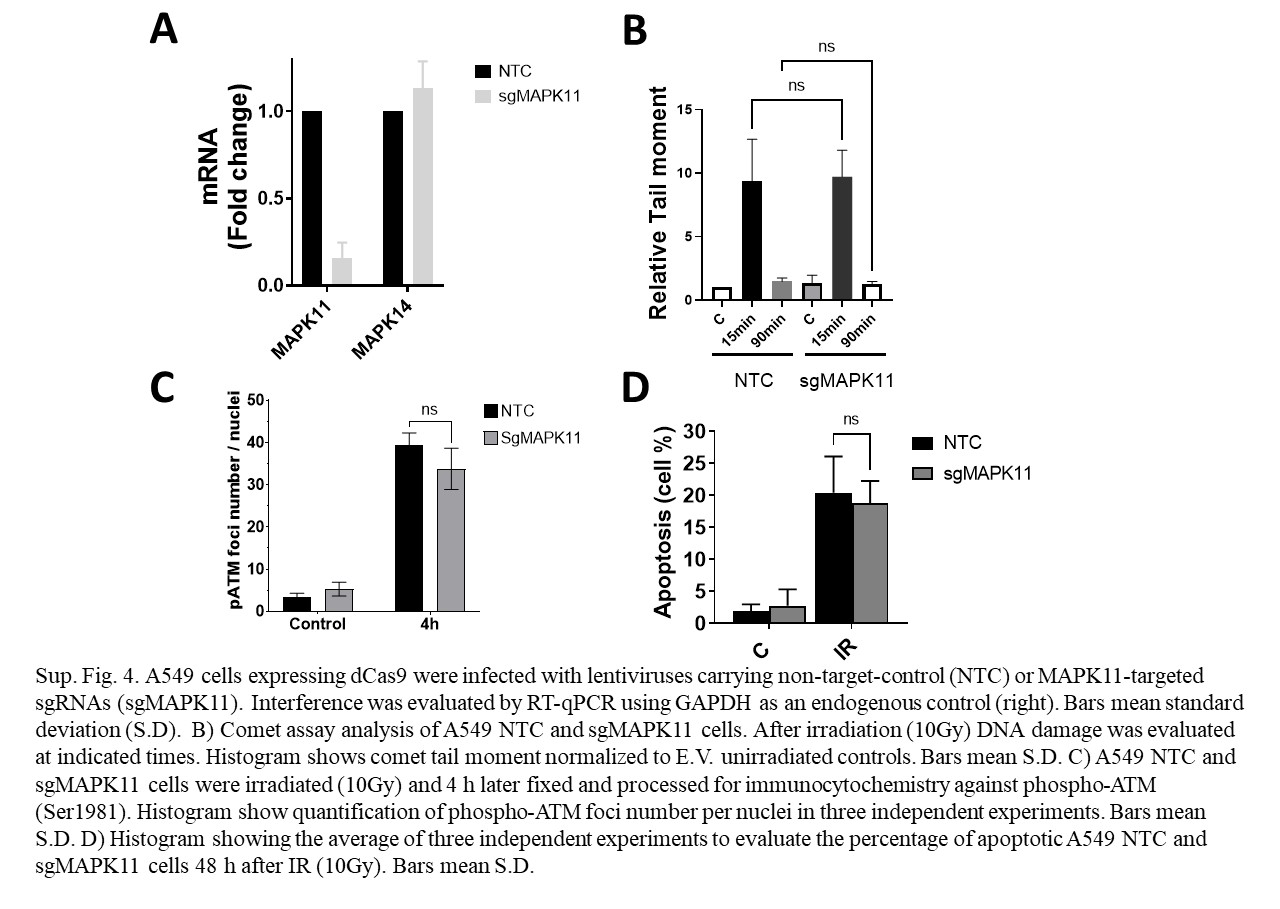

### Supplementary Table 1

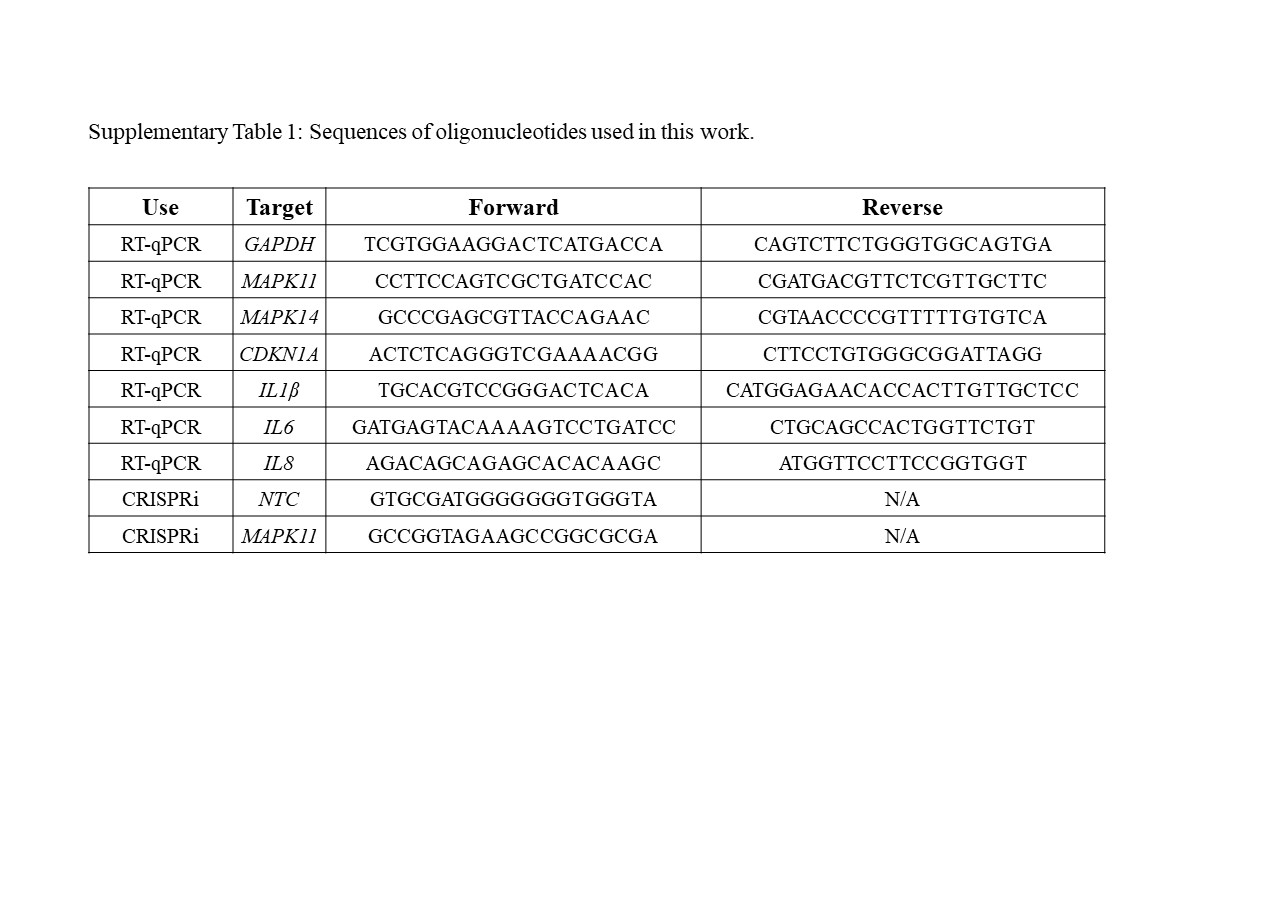

### Supplementary Table 2

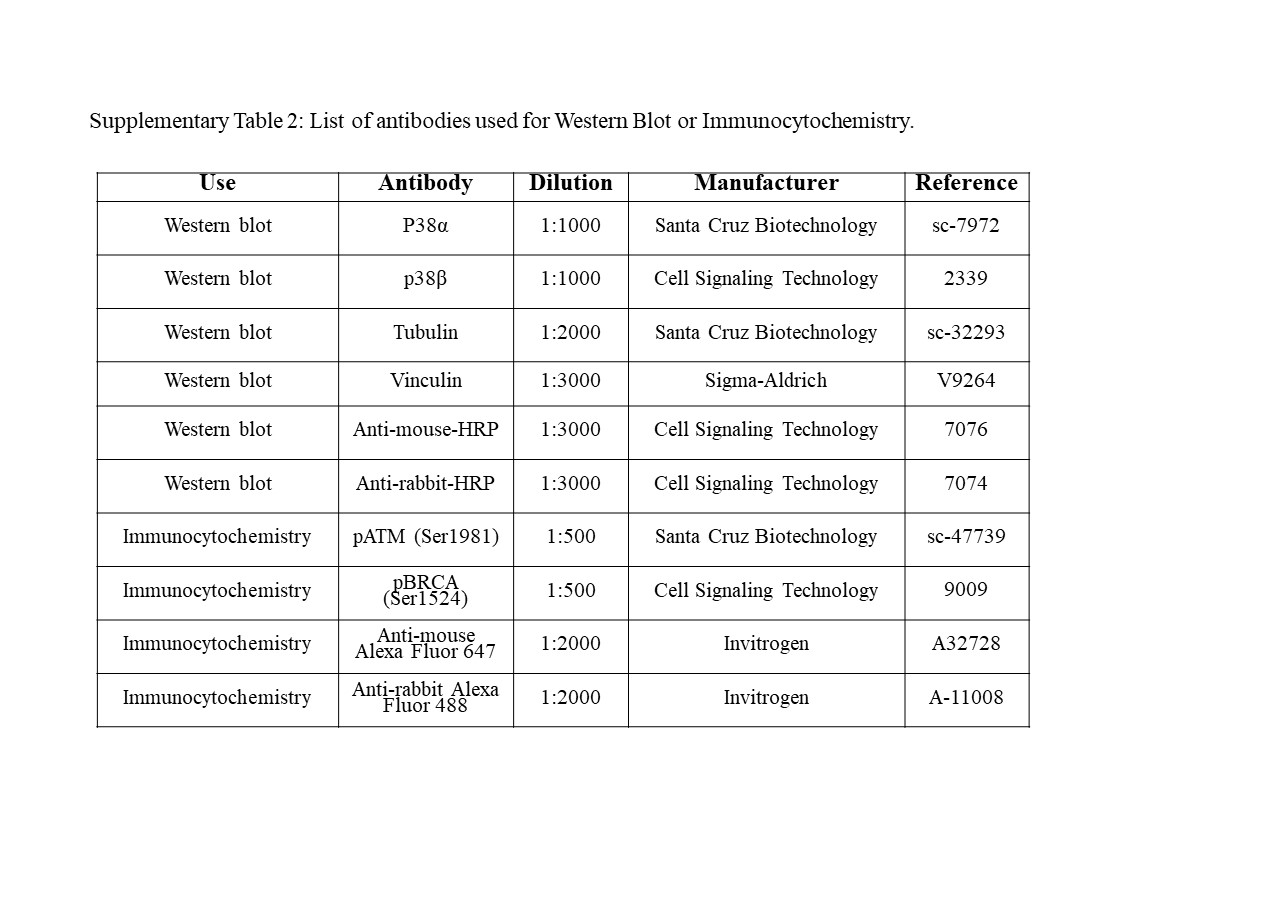
